# Phenotypic plasticity of invasive knotweed across Europe: a distributed common garden experiment

**DOI:** 10.1101/2025.08.18.667133

**Authors:** Ramona Elena Irimia, Madalin Parepa, Nicole Sebesta, Elena Barni, Elisa Giaccone, Yaolin Guo, Sophie Karrenberg, Christina Richards, Oliver Bossdorf

## Abstract

1. Biological invasions expose populations of introduced species to strong selection forces in abiotic and biotic factors, which is expected to lead to adaptive differentiation of populations, particularly for large-scale invaders.
2. To better understand population differentiation and phenotypic plasticity of invasive Japanese knotweed (*Reynoutria japonica*) in Europe, we compared the performance of 45 European knotweed populations, collected across a 2000 km latitudinal transect, in three common gardens with contrasting climatic conditions, one at the southern edge, one in the center, and one at the northern edge of the species’ European distribution.
3. The plants exhibited strong phenotypic plasticity across the three gardens, with a more acquisitive growth strategy in the southern garden, and a more conservative strategy and change of architecture in the climatically unfavorable north. Although we observed phenotypic selection on several leaf traits, and on the plasticity of leaf thickness, with some selection differences between the gardens, there was little evidence of population differentiation, and none for local adaptation. Only for plant architecture (shrubbiness), we detected heritable population variation in trait means and plasticity. However, variation in plasticity was unrelated to climatic variability of origin, and to plant fitness.
4. *Synthesis*. Our results suggest that evolutionary processes - local adaptation and evolution of plasticity - did not play a key role in the success of Japanese knotweed in Europe. Instead, its high baseline plasticity appears to be the key factor that makes it such a strong invader across a broad range of environments.

## 1 INTRODUCTION

Invasive species often expand their ranges across large environmental gradients, which can differ from the conditions in their native range (Zachariah Atwater & Barney 2021). Broad-scale invasions are often linked to either high levels of phenotypic plasticity of these species (Richards *et al*. 2008; Gioria *et al*. 2023) or to their ability to rapidly evolve in the introduced ranges (Whitney & Gabler, 2008; Maron *et al*. 2004; Colautti & Lau 2015). It is important to understand the processes underlying successful invasions, for improving predictions of future range expansions and developing effective management strategies (Gioria *et al*. 2023).

One major explanation for successful invasion across environmental gradients is rapid evolution and adaptive population differentiation. At the regional scale, climate is often a primary driver of natural selection and can lead to clinal differentiation in plant growth, phenology and life-history traits (e.g. Colautti *et al*. 2009, 2010; Keller *et al*. 2009; Woods & Sultan 2022). However, in many introduced species the evolutionary responses to selection may be constrained by the amount of standing genetic variation in phenotype (Colautti *et al*. 2010), with colonization history and founder events expected to leave a major signature on large-scale genetic patterns (Keller *et al*. 2009). In general, local or regional adaptation, resulting from spatially heterogeneous selection is evidenced when local or regional genotypes exhibit higher fitness in their home environment compared to non-local or non-regional genotypes (Kawecki & Ebert 2004).

A key method for investigating phenotypic differentiation among invasive populations, and testing for regional adaptation, are common gardens, where plants of different population origins are grown under standardized conditions, as well as reciprocal transplants among multiple gardens (Clausen *et al*. 1940; Leimu & Fischer 2008; Lortie & Hierro 2021; Schwinning *et al*. 2022). Powerful common garden experiments require the incorporation of a large number of populations sampled across broad environmental gradients, and regional adaptation can only be tested with multiple gardens in sites with contrasting environmental conditions (Colautti & Lau 2015). However, most previous common garden studies of invasive plants did not allow the detection of adaptive differentiation among populations, as they were typically conducted in a single location (but see e.g. Maron *et al*. 2004; Ebeling *et al*. 2011; Colautti & Barrett 2013).

Another important method for understanding geographic patterns of evolution and adaptation are phenotypic selection analyses, where trait variation is related to fitness *in situ* (Lande & Arnold 1983). Such selection analyses can identify contrasting selection pressures in different geographic locations, and thus indicate processes underlying observed evolutionary changes (Siepielski *et al*. 2009; Caruso *et al*. 2017). Although they are important components to understanding adaptive evolution and population differentiation (Colautti & Barrett 2013; Colautti & Lau 2015; Woods & Sultan 2022), selection analyses have remained surprisingly rare in invasion ecology (Colautti & Lau 2015).

Besides adaptive evolution, invasive plants can also adjust to novel environmental conditions through the expression of phenotypic plasticity, and high levels of phenotypic plasticity have been found to be associated with invasion success (Richards *et al*. 2006; Davidson *et al*. 2011; Gioria *et al*. 2021). Moreover, phenotypic plasticity itself is often a heritable variable trait which can rapidly evolve if it offers a fitness advantage in particular environments, such as the introduced ranges of invasive species (Bock *et al*. 2015; Lande 2015). A frequent expectation in invasive plants is that ‘general purpose genotypes’ with a high relative fitness in a wide range of habitats will become successful invaders, or that invasive populations evolve towards greater plasticity (Baker 1965; Richards *et al*. 2008; Wang *et al*. 2025). In general, rapid evolution and phenotypic plasticity may jointly and synergistically contribute to the success of invasive plant populations.

*Reynoutria japonica* Houtt. (Japanese knotweed, Polygonaceae), is a widespread and troublesome invasive plant of temperate Europe and eastern North America. In its introduced range, the species occupies large and highly heterogeneous environments in which it thrives despite strongly reduced genetic diversity (the ‘genetic paradox of invasion’: Allendorf & Lundquist, 2003). In particular, the western European populations of *R. japonica* appear to be composed of clonal descendants of a single introduced genotype, with extremely low genetic diversity (Hollingsworth & Bailey 2000; Zhang *et al*. 2016, 2024). However, a common garden study of 83 *R. japonica* individuals from Central Europe found that these plants, despite their genetic uniformity, displayed differentiation in DNA methylation, as well as in phenotypic traits such as rhizome production, specific leaf area and chlorophyll content, with significant associations between epigenetic and phenotypic variation, and environments of origin (Zhang *et al*. 2016). Recent studies compared trait differences between 50 introduced European populations of *R. japonica* and 58 native populations from China and Japan in two common gardens in China (Cao *et al*. 2024; Wang *et al*. 2025). They found that plants from European populations exhibited increased plasticity in clonality compared to those from the source populations in Kyushu, Japan (Wang *et al*. 2025). Together, these studies indicate that European populations of Japanese knotweed, despite their genetic uniformity, might offer an interesting case for studying the roles of different evolutionary mechanisms, including adaptive phenotypic plasticity, in colonization and invasion of new habitats. However, previous common garden studies took place in the native range, or in a single location in Europe, and they thus did not allow for testing for plasticity, or local adaptation, under invasive-range conditions. These were the goals of our study.

Here, we used a multi-site common garden experiment to study phenotypic differentiation and plasticity of invasive knotweed across Europe. We worked with *R. japonica* from 45 invasive populations spanning a 2000 km latitudinal transect, and we grew these plants in three gardens with contrasting climatic conditions, one at the southern edge, one in the center, and one at the northern edge of the species’ introduced European range. In each garden, we estimated variation in plant performance, as well as in several functional traits, to address the following questions: (1) Do knotweed populations harbor heritable variation in phenotype, and if so, can we find evidence that this is adaptative, i.e. selection favoring different phenotypes in different gardens, or regional populations outperforming more distant ones? (2) How do clonal replicates of the same individual respond to the three garden environments through phenotypic plasticity? (3) Is variation in plasticity related to environmental variability of origin, or to fitness?

## 2 MATERIALS AND METHODS

### 2.1 Study species

Knotweeds are a group of long-lived perennials native to Southeast Asia that were introduced to Europe and North America in the early 19^th^ century as garden ornamentals (Bailey & Conolly 2000). There are three different taxa: *Reynoutria japonica* (Japanese knotweed), *R*. *sachalinensis* (F. Schmidt) Nakai (Giant knotweed) and their hybrid *R*. x *bohemica* Chrtek & Chrtková (Bohemian knotweed). While *Reynoutria japonica* and the hybrid have been spreading aggressively across Europe particularly during the last decades, and are listed as highly invasive, *R*. *sachalinensis* has a sparser distribution and is of less concern (Bailey & Conolly 2000). In its introduced range, the species complex is mostly associated with lowland riparian habitats and spreads primarily vegetatively (Beerling *et al*. 1995).

### 2.2 Plant material

We originally collected knotweed rhizomes in 50 invasive populations in Europe (Table S1), across a 2000 km latitudinal transect from 44°N (Northern Italy) to 59°N (Central Sweden) during the summer of 2019. Our transect spanned most of the species’ latitudinal distribution in Europe, and a substantial climatic gradient, from hot summers and mild winters in Italy to much longer and colder winters in Central Sweden (Table S1). At each field site, we selected five knotweed shoots at regular intervals along a 30 m transect and dug up at least 50 cm rhizome from each (further details about the field sampling can be found in Irimia *et al*. 2025). We determined the taxonomic identity of all individuals and excluded those from five populations that we identified as either *R*. *× bohemica*, or with a mixture of both taxa (Table S1). Consequently, we restricted our analyses to *R*. *japonica* (45 populations). Prior to the set-up of the common garden experiments, we propagated all 225 individuals (45 populations x 5 rhizomes) for two years in the greenhouse. In the spring of 2021, we cut three similar-sized rhizome pieces, each with at least two intact nodes, from each plant and shipped one replicate to each of the three common garden locations.

### 2.3 Common gardens

To investigate phenotypic variation among different European knotweed origins, as well as their phenotypic plasticity, we established three outdoor common gardens with identical plant material and standardized experimental designs in three climatically contrasting locations across Europe (Fig. 1, Table S2). One garden was located near the species’ southern range limit (Botanical garden of the University of Turin; 45.05°N, 7.68°E, 235 m a.s.l), one in the center of the range (Experimental garden at the University of Tübingen; 48.54°N, 9.04°E, 480 m a.s.l) and one near the current northern range limit (Experimental garden at Uppsala University; 59.82°N, 17.65°E, 15 m a.s.l.). In each garden, the experiment had a randomized block design with five spatial blocks (each measuring 11 m x 1.5 m; with 1.5 m distance between blocks). Each block included one randomly selected individual from each of the 45 populations, resulting in a total of 225 plants per garden (5 blocks x 45 individuals), and 675 plants across all three gardens.

**Figure 1.**
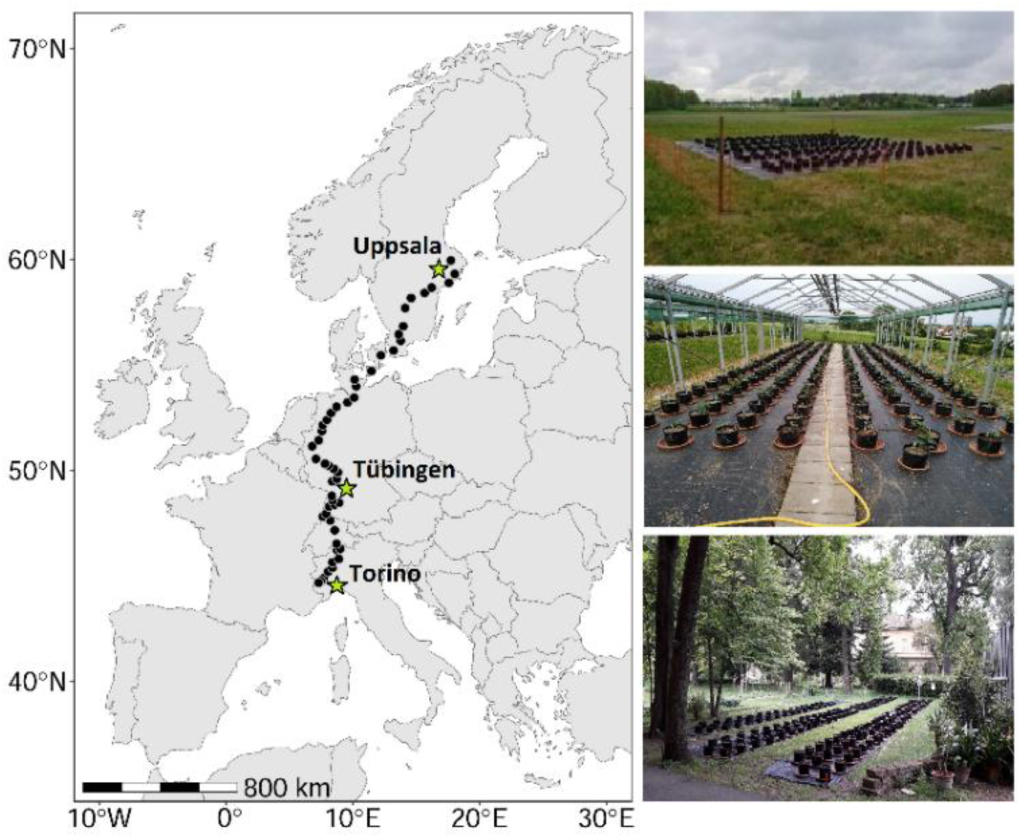
The 50 origins of invasive knotweeds (filled black circles in map), and photos of the three common gardens in Uppsala (top), Tübingen (middle) and Torino (bottom). The green stars on the map indicate the locations of the common gardens.

In all gardens, we planted each rhizome individually into 10 L pots filled with the same commercial potting soil (Patzer Einheitserde Topfsubstrat Type 1; Patzer Erden GmbH, Sinntal, Germany; 340mg N/L, 260 mg P_2_O_5_/L, 330 mg K_2_O/L, 100 mg Mg/L). We planted rhizome fragments horizontally in the center of the pots, at approximately 5-10 cm depth. In each block, we arranged the pots in three rows, with ca. 35 cm distance between pots. All rhizomes were planted in early May 2021 in Torino and Tübingen, and in late May in Uppsala. We grew the plants from May 2021 to June 2022, and fertilized them twice, once in summer 2021 and once in spring 2022, each time with two pellets of long-term fertilizer (Osmocote Exact Protect, 5-6M, 14-8-11 + 2MgO + trace elements; Everris International B.V., Heerlen, Netherlands). In each garden, we watered the plants only when more than 1/3 showed strong signs of wilting (this could range from two to three or four times a month per garden, from May to September). In November 2021, we clipped all aboveground plant biomass and covered the pots with protective fleece for overwintering.

### 2.4 Data collection

As soon as the plants started to re-sprout in spring 2022, we recorded weekly the numbers of shoots taller than 1 cm in each pot. Plants that had not emerged four weeks after the first census were declared dead. In June 2022, we counted the total numbers of shoots in each pot (our measure of vegetative reproduction). We also measured the height and diameter of the tallest shoot in each pot and counted the number of branch tips on this shoot. We then collected five fully developed leaves and measured the leaf chlorophyll content (Minolta SPAD-502, Spectrum Technologies, Inc., Plainfield, IL, USA, expressed in µg/cm^-^²), leaf thickness (Series 293 digital micrometer, Mitutoyo, Tokyo, Japan, expressed in μm), leaf toughness (analog force gauge with 8 mm diameter flat head, Sauter, Balingen, Germany, expressed in J/m^2^) and specific leaf area (SLA; calculated as total leaf area mm^2^ divided by leaf dry mass in mg). We oven-dried the leaf samples at 60°C for 48 h, and subsequently weighed them. We determined leaf area using the software ImageJ (Schneider *et al*. 2012). We also estimated the percent herbivore damage and recorded the presence of pathogens on each plant. Lastly, we attempted to harvest the belowground parts – roots and rhizomes – but because of an extremely dense, entangled fine-root network, we were unable to consistently separate the plant parts, and we did not pursue this further.

### 2.5 Data analyses

We used R v.3.5.1 (R Core Team, 2018) with the *lme4* package (Bates *et al*. 2015) for all analyses, and analyzed seven response variables: the four-leaf traits (chlorophyll content, leaf thickness, leaf toughness and SLA), two growth traits (shoot numbers and (largest) shoot volume), and a plant architecture trait. Shoot volume (expressed in cm^3^) was estimated by the formula:

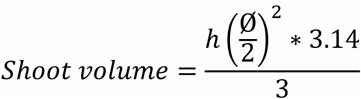

where *h* and Ø were the height and diameter, respectively, of the largest shoot in a pot. The plant architecture variable was defined as the ratio between the number of tips and the stem volume of the largest shoot, which captured variation in lateral branching, i.e. plant shrubbiness.

As a first step, we used the combined data set to explore trait variation among gardens as well as population differences in trait means and plasticities, using generalized linear mixed effects models (GLMMs) with garden, population, and their interaction as fixed factors, and block as a random factor. We assessed the significance of the fixed effects through Type III ANOVA with the Satterthwaite’s method using the *lmerTest* package (Kuznetsova *et al*. 2017). When the model indicated significant garden differences, we applied a Tukey’s contrast with FDR-corrected *P*-values using the *glht()* function in the *multcomp* package (Hothorn *et al*. 2008) to infer which specific gardens differed. To meet the assumptions of a Gaussian distribution of the residuals, the SLA, shoot volume and shrubbiness data were log-transformed prior to the analyses, while the numbers of shoots data were square-root transformed. To visualize differences in plant traits among gardens we also conducted a standardized and centered principal component analysis, using the *prcomp*() function in the package *factoextra* (Kassambara & Mundt 2020), and to further understand the directions of changes across the three gardens, as well as the extent of population variation, we plotted the population-level reaction norms for each response variable.

To explore the association of the leaf traits to plant fitness, we carried out phenotypic selection analyses (Lande & Arnold 1983) that regressed vegetative reproduction (shoot number) or plant growth (shoot volume) on standardized phenotypic traits separately for each garden. To account for environmental variability in the gardens, as well as for the lack of independence of plants from the same population, we included experimental block and population of origin as random factors in the model. We standardized the two fitness-related variables and all phenotypic variables (= leaf traits) to a mean of 0 and a variance of 1, and then calculated relative fitness of individuals by dividing their shoot number or shoot volume by the garden-specific mean of the respective fitness variable. We then calculated both selection differentials (*s*; total selection) as the covariance between relative fitness and each standardized phenotypic trait, and selection gradients (*ß*, direct selection only) as the partial regression coefficient from the multiple regression of relative fitness on all standardized traits (Haggerty & Galloway 2011).

To test for local adaptation of knotweed populations, we calculated the geographic distances between the sites of origin and the sites of the gardens for each population and each garden and used GLMMs to model relative fitness (based on shoot numbers or shoot volumes) as a function of geographic distance, garden and their interaction, with block and population included as random factors (Raabová *et al*. 2007).

Finally, we tested whether variation in phenotypic plasticity was related to fitness, or to the climate variability of population origins. For each trait, we estimated the plasticity of each population through the coefficient of variation (CV) of the mean trait values in the three gardens (Valladares *et al*. 2006). We estimated the across-environment fitness of each population as the mean shoot number or mean volume of the largest shoot across the three gardens, and we used long-term climate data from the Climatic Research Unit (1919-2019; CRU TS4.08, http://doi.org/10/gcmcz3, Harris *et al*. 2024) to calculate inter-annual coefficients of variation for temperature and precipitation at each population origin. We then used linear regression to test for relationships between trait plasticity and plant fitness, and between trait plasticity and climate variability.

## 3 RESULTS

Despite the large climatic differences between the three experimental gardens, the over-winter plant mortality remained low in all, with 4.4% in Torino, 2.2 % in Tübingen and 9.2% in Uppsala. In Torino and Tübingen, most of the plants flowered in 2021 (89% and 63%, respectively), but we did not observe any flowering in Uppsala. There was little to no herbivory and only very few visible leaf pathogens at all three gardens in both years.

### 3.1 Trait differences among gardens and populations

The garden location had a strong effect on all phenotypic variables (Table 1, Table S3). Compared to plants in Tübingen and Torino, the plants in Uppsala produced considerably fewer shoots, and their largest shoots had much smaller volumes, but they were shrubbier and had thicker leaves than the plants in Tübingen and Torino (Fig. 2). In Torino, on the other hand, the plants had consistently higher leaf chlorophyll content and specific leaf area than in Tübingen and Uppsala (Fig. 2). Finally, the plants in Tübingen had higher average leaf toughness values than the plants in the other two gardens.

**Figure 2.**
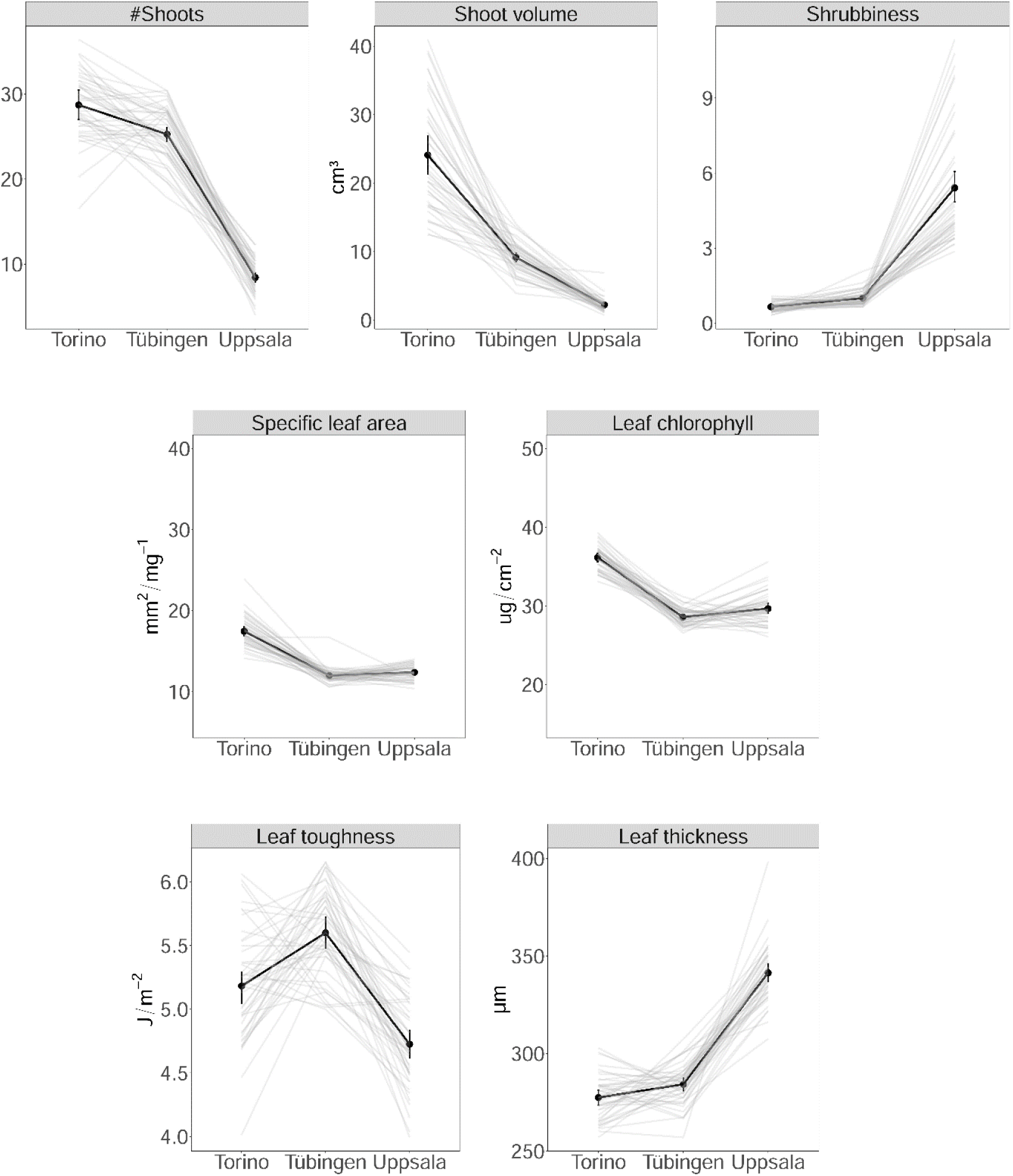
Reaction norms of 45 *R*. *japonica* populations to the three garden environments (in grey) and the grand means (with 95% CI) per garden (in black).

**Table 1.** Effects of garden, population of origin, and their interaction, on growth and leaf traits of *Reynoutria japonica*. The values are *F*-statistics with their corresponding significance levels.

|  |  | Shoot |  |  |  | Leaf | Leaf | Leaf |
| --- | --- | --- | --- | --- | --- | --- | --- | --- |
|  | d.f. | # Shoots | volume | Shrubiness | SLA | chlorophyll | toughness | thickness |
| Garden (G) | 2 | 21.54*** | 53.07*** | 149.0*** | 78.21*** | 30.37*** | 8.112** | 20.95*** |
| Population (P) | 45 | 1.157 | 1.021 | 1.397* | 0.663 | 0.877 | 0.815 | 0.752 |
| P × G | 89 | 0.924 | 1.113 | 1.332* | 0.994 | 0.945 | 0.948 | 1.128 |

d.f. = degrees of freedom. \* $P < 0.05$ , \*\* $P < 0.01$ , \*\*\* $P < 0.001$ .

Besides the garden differences, we found significant population variation in both the mean and the plasticity of plant shrubbiness (Table 1), but in none of the other phenotypic variables. The PCA confirmed distinct trait syndromes particularly for the Torino versus Uppsala gardens, with lower performance and a more conservative strategy – shrubbier growth with thicker leaves – in Uppsala, a more acquisitive strategy with vigorous growth, high chlorophyll and SLA in Torino, and an intermediate position, with less variability, for the Tübingen garden (Fig. 3).

**Figure 3.**
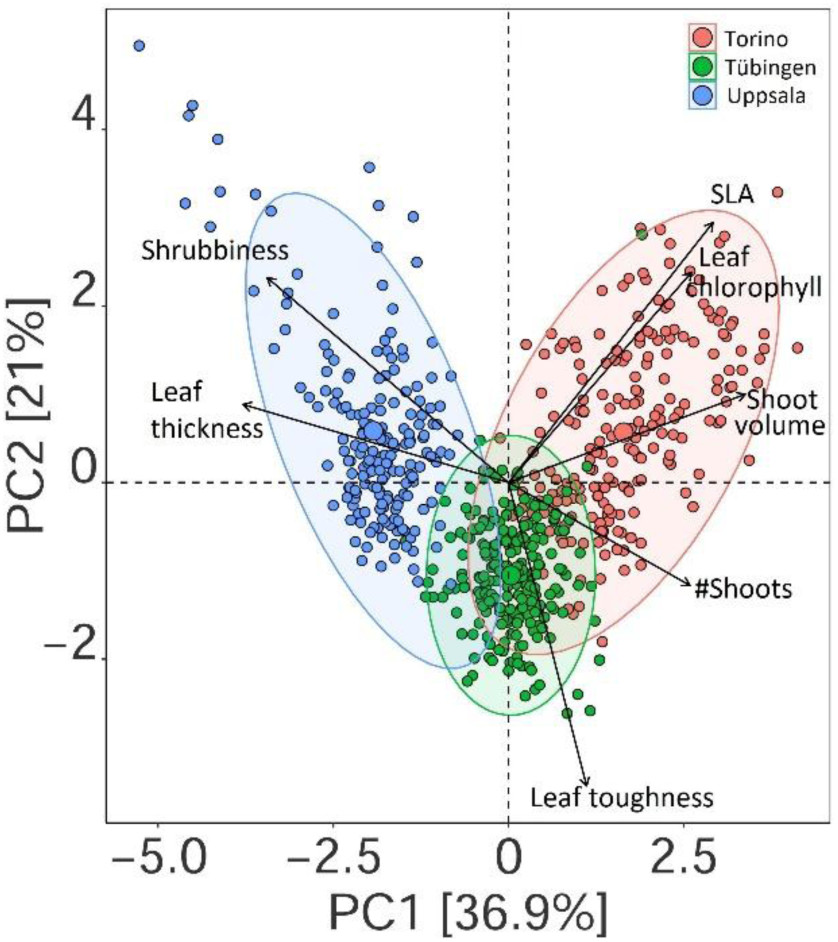
Biplot of the first two principal components, visualizing trait variation in *Reynoutria japonica* across three common gardens in Uppsala (northern range limit), Tübingen (range center) and Torino (southern range limit). Each symbol is an individual plant; the three large symbols indicate the mean values of the gardens, and the shaded ellipses their 95% confidence areas.

### 3.2 Phenotypic selection analysis

Using the number of shoots as a fitness proxy, we found leaf thickness to be positively correlated with fitness in the Torino garden, but negatively correlated with fitness in the Uppsala garden. These patterns were consistent for both total selection (selection differentials) and when only direct selection (selection gradients) was considered (Table 2). We found positive selection on SLA and negative selection on leaf chlorophyll in Uppsala, but no evidence at all for selection based on shoots numbers in the Tübingen garden. On the contrary, using shoot volume as a fitness proxy, we found that patterns of selection were generally negative (Table 2). There was negative selection on SLA in the Torino and Uppsala gardens, on leaf chlorophyll in Tübingen and Uppsala, and on leaf thickness in Torino and Tübingen. Across both fitness proxies, patterns of selection were strongest in the Uppsala garden (7/16 significant tests vs. 4/16 in Torino and 3/16 in Tübingen), and leaf thickness was the trait with the most (7/14) significant fitness associations.

**Table 2.** Standardized linear selection differentials (*s*; total selection) and selection gradients (*ß*; direct selection only) relating leaf trait variation in *Reynoutria japonica* to two measures of plant fitness, when growing in three common garden locations. Significant values are in bold, with FDR corrected *P-*values.

|  |  | # Shoots |  |  |  | Shoot volume |  |  |  |
| --- | --- | --- | --- | --- | --- | --- | --- | --- | --- |
| | | $s$ | $P$-level | $\beta$ | $P$-level | $s$ | $P$-level | $\beta$ | $P$-level |
| Torino | SLA | 0.01 | 0.98 | 0.00 | 0.98 | **-0.16** | **0.04** | -0.07 | 0.37 |
|  | Leaf chlorophyll | 0.03 | 0.84 | 0.02 | 0.98 | 0.07 | 0.32 | 0.06 | 0.66 |
|  | Leaf thickness | **0.24** | **<.001** | **0.23** | **<.001** | **-0.18** | **0.04** | -0.15 | 0.08 |
|  | Leaf toughness | -0.09 | 0.22 | -0.08 | 0.16 | -0.09 | 0.26 | -0.01 | 0.97 |
| Tübingen | SLA | 0.00 | 0.99 | 0.01 | 0.99 | -0.08 | 0.30 | -0.06 | 0.82 |
|  | Leaf chlorophyll | 0.08 | 0.69 | 0.06 | 0.81 | -0.13 | 0.15 | **-0.16** | **0.04** |
|  | Leaf thickness | 0.02 | 0.99 | 0.02 | 0.99 | **-0.23** | **<0.01** | **-0.24** | **<.001** |
|  | Leaf toughness | -0.09 | 0.69 | -0.07 | 0.63 | -0.09 | 0.29 | -0.14 | 0.07 |
| Uppsala | SLA | 0.13 | 0.33 | **0.27** | **<.001** | **-0.29** | **0.01** | **-0.27** | **<.001** |
|  | Leaf chlorophyll | -0.04 | 0.99 | **-0.24** | **<.001** | **-0.21** | **0.04** | -0.14 | 0.06 |
|  | Leaf thickness | **-0.29** | **0.01** | **-0.38** | **<.001** | 0.05 | 0.99 | 0.07 | 0.38 |
|  | Leaf toughness | 0.01 | 0.99 | -0.09 | 0.22 | -0.01 | 0.99 | 0.13 | 0.08 |

### 3.3 Regional adaptation

We found no indication of regional adaptation, i.e. a ‘local *vs*. foreign’ effect where relative fitness decreased with increasing geographic distance (Table S4). There was no relationship between geographic distance and the numbers of shoots produced (*F_1,2_* = 0.19, *P* = 0.66), and there was a *positive* relationship between geographic distance and the volume of the largest shoots (*F_1,2_* = 11.3, *P* < 0.001), i.e. across all three gardens, plants of more distant origins tended to grow more vigorously than plants of regional origin.

### 3.4 Relationships between phenotypic plasticity, fitness and climate variability

We found little evidence for a relationship between the plasticity of individuals from different knotweed populations (CV across the three gardens) and their average, across-garden fitness (Table S5). Only for the plasticity of leaf thickness was there a significant positive relationship (*F*_1,42_ = 6.23, *P* = 0.01) with the average shoot volume. We found no relationship at all between the degree of phenotypic plasticity of plants from different populations and the climatic variability of their origins (Table S6).

## 4 DISCUSSION

Plants may be able to invade broad geographical and environmental ranges because of high levels of phenotypic plasticity, or because they have rapidly adapted to different regions and habitats. To better understand the invasion of Japanese knotweed in Europe, we grew clonal offspring from the entire 2000-km latitudinal European range of the species in three common gardens with contrasting climates, which allowed us to test for plasticity and at the same time trait differentiation, and to examine patterns of selection and regional adaptation.

### 4.1 Phenotypic variation and population differentiation

Overall, plants from the different knotweed populations exhibited very different phenotypes across the three gardens, with distinct trait syndromes between the two range-edge gardens in Torino and Uppsala. In the favorable conditions of the southernmost site—characterized by a warmer climate— the plants displayed a fast resource-use strategy, with enhanced performance through investment in rapid growth and clonality, higher specific leaf area (SLA), and increased leaf chlorophyll content. Under the northern conditions of the Uppsala garden, where low temperatures and shorter day length significantly shorten the growth period, the plants exhibited a much more conservative strategy. This included architectural adjustments and resource allocation toward lateral shoot branching, thicker leaves, and reduced stature. In the intermediate garden in Tübingen the plants displayed trait values that fell between those observed at the two extremes for nearly all measured traits.

Despite the strong general ability of knotweed to express different phenotypes, we found no evidence for population differentiation in traits except for one: plant shrubbiness. These extremely low levels of heritable variation among European populations somewhat contrast with a previous study by Zhang *et al*. (2016) who reported significant regional or population-level variation among 83 Central European origins of Japanese knotweed in multiple traits, including specific leaf area, leaf chlorophyll content, and rhizome production. The study not only analyzed similar traits, but it also cultivated plants for multiple seasons – after several years of pre-cultivation. On the other hand, results from a single common garden should be interpreted with caution (Colautti & Lau 2015), and in Zhang *et al*. (2016) the observed effect sizes were rather small, despite the large geographic range included. We believe our study provides a more complete picture, and that there is little heritable variation among European populations, at least in the leaf traits we measured.

### 4.2 Selection on phenotypic traits

One possible explanation for the absence of population differentiation could be that there are few differences in natural selection experienced by the plants. To get an idea of the potential divergence in selection between contrasting environments we used the leaf traits data from our common gardens, together with the growth data as fitness proxy, to estimate direct and total selection acting on leaf functional traits. We generally found the strongest patterns of selection, i.e. correlations between trait variation and fitness variation, in leaf thickness, followed by SLA and leaf chlorophyll. There were several changes in the direction of selection between gardens, e.g. for selection for increased leaf thickness in Torino but decreased leaf thickness in Tübingen and Uppsala. Moreover, we observed selection for increased SLA in Uppsala, but (weaker) negative selection for it in Torino. Our results thus indicate that there is divergent natural selection on knotweed leaf traits in the three regions – as is generally expected for functional leaf traits along environmental gradients (Wright *et al*. 2004) – and that the lack of population differentiation cannot be explained by a lack of selection differences.

Surprisingly, some of the observed selection was in the opposite direction to the average phenotypes in the three common gardens. For instance, while plants generally had thicker leaves in Uppsala, the selection analyses indicated a fitness advantage to plants with *lower* leaf thickness in Uppsala. Similarly, while plants in Torino generally produced leaves with a larger SLA than those in the other gardens, some selection analyses indicated a fitness advantage of individuals with *lower* SLA. We can only speculate about the causes of these discrepancies. Perhaps our fitness proxies did not capture true fitness well enough, e.g. because they did not include belowground and in particular rhizome biomass (although we attempted to estimate vegetative reproduction through the number of shoots). Another possibility is that the plasticity was maladaptive, e.g. because it evolved under past conditions, or it incurred significant costs (Murren *et al*. 2015). Alternatively, the selection results may be misleading if they reflect indirect selection (because of genetic correlations). Ultimately, only more in-depth studies with improved fitness measures, and following the performance of different phenotypes over longer timescales will be able to answer these questions.

### 4.3 Regional adaptation

We found no evidence for regional adaptation of European *R*. *japonica* populations in this study. The performance of plants was not higher in populations originating closer to the common gardens as compared to those originating from larger distances (Raabová *et al*. 2007). This lack of regional adaptation is not surprising, given the little population differentiation we detected, which is presumably a consequence of the largely clonal spread of the species in Europe. *Reynoutria japonica* was introduced to Europe in the mid-19^th^ century and spread rapidly across the continent through horticultural trade, which was based on rhizome fragments (Bailey & Conolly 2000). A previous study reported that the same individuals from across the 45 European populations all belonged to the same haplotype that is found in the putative source populations in Kyushu (Zhang *et al*. 2024). While Zhang *et al*. (2016) observed higher levels of epigenetic and phenotypic variation in nearby European populations, our study could not confirm these findings. The situation is slightly different in the North American invasive range which has a more complex introduction history, with slightly more genetic diversity among populations (Del Tredici 2017). A reciprocal transplant between three *R*. *japonica* sites in the northern, central and southern parts, respectively, of the species’ US range found weak support for regional adaptation in clonality but only in the northern site (Van Wallendael *et al*. 2018). Another reciprocal transplant between beach, marsh and roadside sites on Long Island (New York), Yuan *et al*. (2024) also found mixed evidence for local adaptation, with greater biomass of marsh plants, and greater survival of beach and roadside plants, in their respective home sites, compared to plants from other habitat origins. Therefore, the overall evidence for local or regional adaptation of *R. japonica* remains weak, similar to many other clonal plant invaders that may demonstrate less local adaptation than invasive plants in general (Oduor *et al*. 2016; but see: Lenssen *et al*. 2004; Simón-Porcar *et al*. 2021).

### 4.4 Phenotypic plasticity and evolutionary potential

Across all populations of origin, Japanese knotweed showed strong plastic responses to the environmental conditions of the three gardens, with significant changes in all measured traits, and an overall response characteristic of a *’jack-and-master*’ strategy (sensu Richards *et al*. 2006). The species appears to be able to maintain growth in a broad range of conditions, but at the same time to take advantage of more favorable conditions – a combination of robustness and opportunism. Despite this strong overall phenotypic plasticity, we found little evidence of genetic variation in plasticity. Only with regard to plant architecture (shrubbiness) was there a significant origin by environment interaction. Architectural plasticity is known to be a key trait for plants to improve resource acquisition under different resource conditions (Pierik *et al*. 2021). A recent experimental study by Martin *et al*. (2020) also found *Reynoutria japonica* to strongly respond to habitat quality – in this case shade and mowing – by plasticity of architectural traits such as lateral spread, shoot density, and clonal integration. Under stressful conditions, the species adopted a very different growth strategy from that observed in favorable environments (e.g. full light), with more widely spaced horizontally spreading ramets, suggesting an escape or foraging strategy (Martin *et al*. 2020). Notably, we also observed this morphotype in our common garden in Uppsala.

We found little evidence for adaptive variation in plasticity among individuals. Only for leaf thickness was there a significant positive association between the average plasticity of individuals and their across-environment fitness proxy (plant size). It is possible that enhanced leaf mesophyll density and leaf thickening enhance the drought resistance of Japanese knotweed. However, further research is needed to understand this relationship better, and to explore potential trade-offs between plasticity and different fitness components, particularly in investment in below-ground biomass and rhizome production.

## 5 CONCLUSIONS

Studying the phenotypic variation and plasticity of invasive species in their introduced ranges is essential to understand the invasion dynamics and success of these species. The results of our multi- common garden experiment demonstrate that *Reynoutria japonica* populations in Europe have a strong capacity for phenotypic plasticity, which allows them to adjust their growth strategies to a broad range of climatic conditions. As we found little evidence of population differentiation in trait means or plasticity, our results also indicate that evolutionary processes - local adaptation and evolution of plasticity - did not play a significant role in the success of Japanese knotweed in Europe. Instead, its high baseline plasticity may be the key factor that makes it such a strong invader across a broad range of environments.

