## supplements for "Phenotypic plasticity of invasive knotweed across Europe: a distributed common garden experiment"

**SUPPLEMENTARY MATERIAL**

**Table S1.** Geographic locations of the 50 Japanese knotweed s.l. populations, sampled across a latitudinal gradient in the introduced European range, from northern Italy to central Sweden, with the long-term average (CRU 1919-2019) annual temperature and precipitation values for each. We excluded from the analyses the five populations that we identified as either *R.* × *bohemica* or with a mixture of both taxa.

| Country | Population | Latitude | Longitude | Elevation [m] | Ploidy | Taxa | Annual mean T [°C] | Annual P [mm] |
|---|---|---|---|---|---|---|---|---|
| Italy | 1 | 44.8836 | 7.68843 | 230 | 8X | *R. japonica* | 12.7 | 726 |
| Italy | 2 | 44.7475 | 7.47791 | 258 | 8X | *R. japonica* | 8.48 | 988 |
| Italy | 3 | 44.6711 | 7.28597 | 586 | 8X | *R. japonica* | 8.48 | 988 |
| Italy | 4 | 45.1911 | 8.03850 | 159 | 8X | *R. japonica* | 12.4 | 836 |
| Italy | 5 | 45.3811 | 8.37205 | 135 | 6X | *R.* × *bohemica* | 12.0 | 920 |
| Italy | 6 | 45.6365 | 8.37471 | 308 | 6X, 8X | mixed | 9.18 | 1132 |
| Italy | 7 | 45.7998 | 8.87095 | 336 | 8X | *R. japonica* | 11.3 | 1219 |
| Switzerland | 8 | 46.2133 | 8.74464 | 374 | 6X | *R.* × *bohemica* | 9.40 | 1286 |
| Switzerland | 9 | 46.2730 | 8.99511 | 258 | 8X | *R. japonica* | 6.80 | 1411 |
| Switzerland | 10 | 46.5135 | 8.69992 | 1003 | 8X | *R. japonica* | 2.70 | 1549 |
| Switzerland | 11 | 47.1553 | 8.56083 | 720 | 8X | *R. japonica* | 8.20 | 1155 |
| Germany | 12 | 47.6240 | 8.21540 | 344 | 8X | *R. japonica* | 7.70 | 1087 |
| Germany | 13 | 47.8119 | 7.61194 | 240 | 8X | *R. japonica* | 8.90 | 914 |
| Germany | 14 | 47.9401 | 7.87734 | 633 | 8X | *R. japonica* | 8.90 | 914 |
| Germany | 15 | 48.2796 | 8.11127 | 298 | 8X | *R. japonica* | 7.90 | 980 |
| Germany | 16 | 48.3737 | 8.55788 | 575 | 8X | *R. japonica* | 8.10 | 934 |
| Germany | 17 | 48.4655 | 8.91653 | 419 | 8X | *R. japonica* | 8.10 | 934 |
| Germany | 18 | 48.5637 | 8.39383 | 513 | 8X | *R. japonica* | 9.50 | 814 |
| Germany | 19 | 48.7851 | 8.32216 | 193 | 8X | *R. japonica* | 9.50 | 814 |
| Germany | 20 | 49.4768 | 8.36521 | 95 | 6X | *R.* × *bohemica* | 10.1 | 630 |
| Germany | 21 | 49.5892 | 8.73014 | 169 | 8X | *R. japonica* | 9.50 | 663 |
| Germany | 22 | 49.9048 | 8.83001 | 145 | 8X | *R. japonica* | 9.50 | 663 |

| | | | | | | | | |
|---|---|---|---|---|---|---|---|---|
| Germany | 23 | 50.0706 | 8.48126 | 107 | 8X | *R. japonica* | 9.10 | 650 |
| Germany | 24 | 50.2307 | 8.05309 | 230 | 6X | *R. × bohemica* | 8.60 | 676 |
| Germany | 25 | 50.3141 | 7.78856 | 130 | 8X | *R. japonica* | 9.10 | 678 |
| Germany | 26 | 50.5408 | 7.08143 | 205 | 8X | *R. japonica* | 9.80 | 782 |
| Germany | 27 | 51.1425 | 6.78324 | 42 | 8X | *R. japonica* | 10.3 | 819 |
| Germany | 28 | 51.4272 | 7.29018 | 77 | 8X | *R. japonica* | 9.50 | 862 |
| Germany | 29 | 51.8657 | 7.548340 | 64 | 8X | *R. japonica* | 9.70 | 756 |
| Germany | 30 | 52.1518 | 7.61590 | 40 | 8X | *R. japonica* | 9.40 | 759 |
| Germany | 31 | 52.3888 | 7.94417 | 49 | 8X | *R. japonica* | 9.40 | 759 |
| Germany | 32 | 52.7201 | 8.26118 | 29 | 8X | *R. japonica* | 9.20 | 768 |
| Germany | 33 | 53.0117 | 8.70257 | 8 | 8X | *R. japonica* | 9.10 | 745 |
| Germany | 34 | 53.2206 | 9.56834 | 34 | 8X | *R. japonica* | 8.80 | 739 |
| Germany | 35 | 53.4468 | 10.0835 | 2 | 8X | *R. japonica* | 8.90 | 681 |
| Germany | 36 | 53.9850 | 10.2483 | 39 | 8X | *R. japonica* | 8.80 | 724 |
| Germany | 37 | 54.3061 | 10.1328 | 11 | 8X | *R. japonica* | 8.60 | 744 |
| Denmark | 38 | 54.7106 | 11.4508 | 4 | 8X | *R. japonica* | 8.70 | 604 |
| Denmark | 39 | 55.4556 | 12.1882 | -2 | 8X | *R. japonica* | 8.50 | 576 |
| Sweden | 40 | 55.6822 | 13.1864 | 14 | 8X | *R. japonica* | 7.80 | 675 |
| Sweden | 41 | 56.1447 | 13.7575 | 50 | 8X | *R. japonica* | 7.10 | 764 |
| Sweden | 42 | 56.4450 | 13.6029 | 105 | 8X | *R. japonica* | 7.10 | 764 |
| Sweden | 43 | 56.8325 | 13.9603 | 150 | 8X | *R. japonica* | 6.70 | 824 |
| Sweden | 44 | 57.6978 | 14.1110 | 188 | 8X | *R. japonica* | 5.70 | 655 |
| Sweden | 45 | 58.16846 | 14.57616 | 144 | 8X | *R. japonica* | 6.00 | 600 |
| Sweden | 46 | 58.41322 | 15.65144 | 45 | 8X | *R. japonica* | 6.30 | 563 |
| Sweden | 47 | 58.66624 | 16.20078 | 9 | 8X | *R. japonica* | 6.20 | 571 |
| Sweden | 48 | 58.89243 | 17.56211 | 17 | 8X | *R. japonica* | 7.00 | 479 |
| Sweden | 49 | 59.31895 | 18.01697 | 12 | 8X | *R. japonica* | 7.00 | 494 |
| Sweden | 50 | 59.94927 | 17.70784 | 51 | 8X | *R. japonica* | 6.10 | 536 |

5

**Table S2**: The monthly temperature averages, with minimum and maximum temperatures in brackets, as well as total monthly precipitation values at the three common garden locations during the time of our experiment (data from https://en.tutiempo.net/climate).

| Year | Month | Temperature [°C] | | | Precipitation [mm] | | |
|---|---|---|---|---|---|---|---|
| | | Torino | Tübingen | Uppsala | Torino | Tübingen | Uppsala |
| 2021 | May | 15.6 (9.40-21.0) | 11.4 (6.70-16.1) | 9.60 (3.80-14.8) | 73.9 | 68.1 | 71.4 |
| | June | 22.6 (17.2-28.1) | 20.0 (13.8-25.9) | 17.4 (10.5-23.7) | 92.5 | 174 | 60.2 |
| | July | 23.3 (18.1-28.5) | 19.0 (13.7-24.1) | 19.4 (11.5-26.3) | 76.2 | 83.1 | 29.2 |
| | August | 23.1 (17.4-28.6) | 17.3 (12.8-21.9) | 14.6 (9.50-19.3) | 25.9 | 104 | 138 |
| | September | 20.6 (15.6-25.8) | 16.0 (9.80-22.3) | 11.1 (6.00-15.7) | 35.0 | 27.6 | 30.2 |
| | October | 12.9 (8.20-18.0) | 9.50 (4.30-15.5) | 8.40 (4.50-11.8) | 44.0 | 37.1 | 50.5 |
| | November | 7.60 (3.70-11.6) | 3.60 (0.50-6.90) | 1.70 (-2.00-5.00) | 97.3 | 20.3 | 23.6 |
| | December | 3.10 (-1.40-8.80) | 3.80 (1.00-6.60) | -4.00 (-8.20-(-0.5) | 14.5 | 50.0 | 21.6 |
| 2022 | January | 3.10 (-2.70-10.2) | 2.10 (-1.00-5.60) | -0.60 (-3.80-2.20) | 0.80 | 27.2 | 55.6 |
| | February | 6.50 (0.20-13.1) | 5.10 (0.80 -9.30) | -0.40 (-3.70-2.60) | 1.50 | 29.7 | 24.1 |
| | March | 8.40 (2.60-13.5) | 6.00 (-0.50-12.6) | 2.00 (-3.50-8.00) | 4.30 | 22.6 | 0.75 |
| | April | 13.0 (7.10-18.4) | 8.60 (3.40-13.4) | 4.00 (-2.70-10.0) | 23.6 | 95.2 | 33.0 |
| | May | 19.7 (14.4-24.9) | 15.8 (9.00-21.4) | 10.6 (3.80-16.4) | 98.8 | 53.5 | 28.9 |
| | June | 24.4 (18.5-29.7) | 19.8 (13.2-25.6) | 17.0 (10.6-22.6) | 30.5 | 109 | 54.3 |

**Table S3.** Post-hoc pairwise comparisons of different gardens for each of the phenotypic variables measured in *R. japonica*, with *P*-values after FDR correction, and significant comparisons in bold.

| Phenotypic variable | Comparison | Difference of means | SE of differences | 95% CI | z-value | Adjusted *P*-value |
|---|---|---|---|---|---|---|
| # Shoots | Tübingen-Torino | -0.20 | 0.40 | [-1.15, 0.73] | -0.51 | 0.60 |
| | Uppsala-Torino | -2.38 | 0.40 | [-3.32, -1.43] | -5.91 | **<.001** |
| | Uppsala-Tübingen | -2.17 | 0.40 | [-3.11, -1.23] | -5.40 | **<.001** |
| Shoot volume | Tübingen-Torino | -0.30 | 0.09 | [-0.52, -0.08] | -3.20 | **.001** |
| | Uppsala-Torino | -0.98 | 0.09 | [-1.20, -0.75] | -10.2 | **<.001** |
| | Uppsala-Tübingen | -0.67 | 0.09 | [-0.89, -0.44] | -7.03 | **<.001** |
| Shrubbiness | Tübingen-Torino | 0.24 | 0.08 | [0.04, 0.43] | 2.92 | 0.05 |
| | Uppsala-Torino | 0.94 | 0.08 | [0.74, 1.13] | 11.3 | **<.001** |
| | Uppsala-Tübingen | 0.70 | 0.08 | [0.50, 0.89] | 8.39 | **<.001** |
| Leaf chlorophyll | Tübingen-Torino | -7.58 | 1.02 | [-9.98, -5.17] | -7.38 | **<.001** |
| | Uppsala-Torino | -6.45 | 1.03 | [-8.87, -4.03] | -6.24 | **<.001** |
| | Uppsala-Tübingen | 1.12 | 1.03 | [-1.29, 3.54] | 1.09 | 0.27 |
| Leaf thickness | Tübingen-Torino | 7.85 | 10.5 | [-16.9, 32.6] | 0.73 | 0.45 |
| | Uppsala-Torino | 64.2 | 10.6 | [39.4, 89.1] | 6.06 | **<.001** |
| | Uppsala-Tübingen | 56.4 | 10.6 | [31.5, 81.2] | 5.32 | **<.001** |
| Leaf toughness | Tübingen-Torino | 0.41 | 0.21 | [-0.09, 0.92] | 1.88 | 0.05 |
| | Uppsala-Torino | -0.45 | 0.21 | [-0.97, 0.05] | -2.06 | 0.05 |
| | Uppsala-Tübingen | -0.86 | 0.21 | [-1.38, -0.35] | -3.95 | **<.001** |
| SLA | Tübingen-Torino | -0.15 | 0.01 | [-0.19, -0.12] | -10.6 | **<.001** |
| | Uppsala-Torino | -0.14 | 0.01 | [-0.18, -0.11] | -9.65 | **<.001** |
| | Uppsala-Tübingen | 0.01 | 0.01 | [-0.02, 0.04] | 0.87 | 0.38 |

**Table S4.** ANOVA testing the effect of the geographic distance (i.e. 'local vs. foreign' advantage), on *R. japonica* fitness (vegetative reproduction or growth). Significant effects are in bold.

| | | # Shoots | | Shoot volume | |
|---|---|---|---|---|---|
| | d.f. | *F*-ratio | *P*-value | *F*-ratio | *P*-value |
| Garden (G) | 2 | **18.49** | **<.001** | **45.54** | **<.001** |
| Geographic distance (GD) | 1 | 0.19 | 0.66 | **11.37** | **<.001** |
| G × GD | 2 | 0.15 | 0.85 | 0.05 | 0.94 |

**Table S5**. Results of univariate linear regression testing for relationships between trait plasticity (calculated as the coefficient of variation for each trait, across the three gardens) and average fitness across the three gardens (measured as number of shoots and volume of the largest shoot). Shown are *df*-degrees of freedom, *F*-ratios, and *P*-values. Significant results are highlighted in bold.

| | | *# Shoots* | | *Shoot volume* | |
|---|---|---|---|---|---|
| | d.f. | *F*-ratio | *P*-value | *F*-ratio | *P*-value |
| CV of SLA | 44 | 0.09 | 0.75 | 0.00 | 0.99 |
| CV of leaf chlorophyll | 43 | 0.31 | 0.57 | 0.55 | 0.46 |
| CV of leaf thickness | 42 | 3.51 | 0.06. | **6.23** | **0.01** |
| CV of leaf toughness | 41 | 1.70 | 0.19 | 0.82 | 0.36 |

**Table S6**. Results of linear regressions testing for relationships between population-level variation in trait plasticity (coefficient of variation of the population-level trait means in three experimental gardens in Torino, Tübingen and Uppsala) across 45 European populations of *Reynoutria japonica*, and the climate variability of their population origins (inter-annual CV based on 1919-2019 data for annual mean temperature or total annual precipitation).

| | | # Shoots | | Shoot volume | | Shrubbiness | | SLA | | Leaf chlorophyll | | Leaf toughness | | Leaf thickness | |
|---|---|---|---|---|---|---|---|---|---|---|---|---|---|---|---|
| | d.f. | *F*-ratio | *P*-level | *F*-ratio | *P*-level | *F*-ratio | *P*-level | *F*-ratio | *P*-level | *F*-ratio | *P*-level | *F*-ratio | *P*-level | *F*-ratio | *P*-level |
| CV Temperature | 43 | 3.35 | 0.07. | 2.02 | 0.16 | 0.07 | 0.79 | 0.61 | 0.43 | 1.34 | 0.25 | 0.20 | 0.65 | 1.07 | 0.30 |
| CV Precipitation | 42 | 1.93 | 0.17 | 0.16 | 0.68 | 1.40 | 0.24 | 0.20 | 0.65 | 3.62 | 0.06. | 0.19 | 0.66 | 0.04 | 0.82 |
| CV Temp × CV Precip | 41 | 2.77 | 0.10 | 0.67 | 0.41 | 0.98 | 0.32 | 0.03 | 0.85 | 0.46 | 0.49 | 0.01 | 0.89 | 0.27 | 0.59 |
